# Costs and benefits of sub-lethal Drosophila C virus infection

**DOI:** 10.1101/111930

**Authors:** Pedro F. Vale, Vanika Gupta, Charlotte Stewart, Samuel S.C. Rund, Katy Monteith

## Abstract

Viruses are major evolutionary drivers of insect immune systems. Much of our knowledge of insect immune responses derives from experimental infections using the fruit fly *Drosophila melanogaster.* Most experiments, however, employ lethal pathogen doses through septic injury, frequently overwhelming host physiology. While this approach has revealed a number of immune mechanisms, it is less informative about the fitness costs hosts may experience during infection in the wild. Using both systemic and oral infection routes we find that even apparently benign, sub-lethal infections with the horizontally transmitted Drosophila C Virus (DCV) can cause significant physiological and behavioral morbidity that is relevant for host fitness. We describe DCV-induced effects on fly reproductive output, digestive health, and locomotor activity, and we find that viral morbidity varies according to the concentration of pathogen inoculum, host genetic background and sex. Notably, sub-lethal DCV infection resulted in a significant increase in fly reproduction, but this effect depended on host genotype. We discuss the relevance of sub-lethal morbidity for *Drosophila* ecology and evolution, and more broadly, we remark on the implications of deleterious and beneficial infections for the evolution of insect immunity.

## INTRODUCTION

Viral infections are pervasive throughout the living world (Suttle, 2005; Rosario & Breitbart, 2011). Viruses of insects have attracted considerable interest (Miller & Ball, eds, 1998), in part due to their potential role in the bio-control of insect pests (Lacey *et al*., 2015), and also because insects are vectors of many viral pathogens of plants (Whitfield *et al*., 2015), animals and humans (Conway *et al.*, 2014). The abundance and diversity of insect viruses, combined with the extensive morbidity and mortality they cause, make viral infections potentially powerful determinants of insect population dynamics and evolution (Dwyer *et al*., 2004;Obbard *et al.*, 2006;Wilfert *et al.*, 2016).

Much of our knowledge of insect immune responses to viral infections has come from work using the fruit fly *Drosophila melanogaster*, where the focus has been on elucidating the genetics underlying antiviral immunity (Dostert *et al.*, 2005; Huszar & Imler, 2008; Kemp & Imler, 2009; Sabin *et al.*, 2010;Magwire *et al.*, 2012). Several RNA viruses have been described and investigated in this context, including Nora virus (Habayeb *et al.*, 2009), Drosophila A virus (DAV)(Ambrose *et al.*, 2009), Flock House Virus (FHV) (Scotti *et al.*, 1983) and Drosophila C Virus (DCV) (Jousset *et al.*, 1977), a horizontally transmitted ssRNA virus in the *Dicistroviridae* family (Huszar & Imler, 2008). Initial investigations of DCV infection found that it replicates in the fly’s reproductive and digestive tissues (Lautié-Harivel & Thomas-Orillard, 1990) and that infection results in accelerated larval development but also causes mortality (Thomas-Orillard, 1984; Gomariz-Zilber *et al.*, 1995). More recent work has shown that systemic infection with elevated concentrations of DCV causes pathology within the fly’s food storage organ, the crop, leading to intestinal obstruction, lower metabolic rate and reduced locomotor activity (Arnold *et al.*, 2013;Chtarbanova *et al.*, 2014). There is also considerable genetic variation in fly survival when challenged systemically with DCV, which appears to be controlled by few genes of large effect (Magwire *et al.*, 2012).

While this level of detail concerning the physiological consequences and the underlying genetics of infection is remarkable, it is important to recognize that our knowledge of viral infections comes almost entirely from experimental infections that challenge model systems, such as *Drosophila*, with artificially high viral concentrations during systemic infections. Even in cases where natural routes of infection have been investigated (Gomariz-Zilber *et al.*, 1995; Ferreira *et al.*, 2014; Stevanovic & Johnson, 2015; Vale & Jardine, 2015), these have often been achieved by using much higher doses than flies are likely to encounter in the wild in order to cause significant mortality. Highly lethal systemic or oral infections have been useful in unravelling broad antiviral immune mechanisms (Dostert *et al.*, 2005; Wang *et al.*, 2006; Kemp & Imler, 2009; Nayak *et al.*, 2013;Karlikow *et al.*, 2014), but it is unlikely that the morbidity and mortality they cause is an accurate reflection of the level of disease experienced by flies in the wild, where viral infections appear to be widespread among many species of *Drosophila* as low level persistent infections with apparently little pathology (Kapun *et al.*, 2010;Webster *et al.*, 2015). Our understanding of the fitness costs of viral infection in *Drosophila* is therefore severely limited, which is striking given the evidence from population genetic data that viruses are major drivers of adaptive evolution in *Drosophila* immune genes (Obbard *et al.*, 2006, 2009; Early *et al.*, 2016).

To gain a better understanding of the potential fitness costs of DCV infection, we measured the physiological and behavioural responses of flies challenged with either a low, sub-lethal concentration of DCV through the oral route of infection or when exposed to a range of sub-lethal viral concentrations systemically through intra-thoracic injury. We focused on traits that have been previously shown to be affected by DCV infection such as survival, fecal excretion, and locomotor activity, as well as female reproductive output, which is ultimately important for evolutionary fitness. We find that even apparently benign, sub-lethal infections can cause significant physiological and behavioural morbidity that is relevant to fly fitness, and that these effects vary according to viral concentration, host genetic background and sex.

## MATERIAL AND METHODS

### Fly lines and rearing conditions

In experiment 1 (systemic DCV infection) we used *Drosophila melanogaster* line *G9a^+/+^* described previously (Merkling *et al.*, 2015), kindly provided by R. van Rij (Radboud University, Nijmegen, NL). This line was maintained on standard Lewis Cornmeal medium (Lewis, 2014) under standard laboratory conditions at 25°C, 12h: 12h Light:Dark cycle. Experimental flies were generated by setting up 20 replicate Lewis vials with 15 males and 15 females to mate and lay eggs for 24 hours. Three-to-four-day-old adults that eclosed from the eggs laid during this period were infected systemically (see below) and then followed individually for health measures.

In experiment 2 (oral DCV exposure) we used ten *D. melanogaster* lines from the Drosophila Genetic Reference Panel (DGRP): RAL-83, RAL-91, RAL-158, RAL-237, RAL-287, RAL-317, RAL-358, RAL-491, RAL-732, and RAL-821. Given we had no prior knowledge of how the DGRP panel vary in response to oral DCV infection, these lines were chosen randomly. All lines were previously cleared of *Wolbachia* and have been maintained *Wolbachia-free* for at least 3 years. Fly stocks were kept at a density of 30 individuals in bottles on standard Lewis medium at 24.5± 0.5°C. Flies were allowed to mate and lay eggs for three days and then removed. When eggs had developed into three-day old imagoes, we picked 16 male and 16 female flies at random from each DGRP line (320 flies in total). Half of these flies (n=8 replicates) were individually exposed to DCV through the oral route of infection (see details below) and the other half were exposed to a sterile Ringers solution (7.2 g/L NaCl; 0.17 g/L CaCl_2_; 0.37 g/L KCl, diluted in sterile water, pH 7.4) as a control (n=8 replicates). Following infection, all flies were kept individually in vials kept in incubators at 24.5°C ± 0.5 with a 12h:12h light:dark cycle for the remainder of the experiment. Vials were randomized within trays to reduce any positional effects within incubators.

### DCV stock and culturing

The Drosophila C Virus (DCV) isolate used in both experiments was originally isolated in Charolles, France (Jousset *et al.*, 1977), and was produced in Drosophila line 2 (DL2) cells as described previously (Longdon *et al.*, 2013; Vale & Jardine, 2015). Infectivity of the virus was calculated by measuring cytopathic effects in DL2 cells using the Reed-Muench end-point method to calculate the Tissue Culture Infective Dose 50 (TCID_50_) (Reed & Muench, 1938). The DCV stock used in this experiment had an infectivity of approximately 4×10^9^ DCV infectious units (IU)/mL. This stock culture was serially diluted to achieve the desired concentrations (approximately 10^2^ 10^3^ and 10^5^ DCV IU/mL for systemic infection and 10^5^ DCV IU/mL for oral infection) and kept at −80°C until needed.

### Systemic DCV infection and viral titers

We exposed 20 individual male and female flies to each of 4 viral concentrations (160 flies in total)− 0 (control), 10^2^, 10^3^ and 10^5^ DCV IU/ml, obtained by serial diluting the viral stock with 10mM Tris-HCl (pH 7.3). Flies were infected systemically by intra-thoracic pricking with a needle immersed in DCV suspension under light CO_2_ anesthesia. Control flies were pricked with a needle dipped in sterile10mM Tris-HCl (pH 7.3). An additional five individuals for each sex/dose combination were infected as described above to quantify DCV within flies following infection, using the expression of DCV RNA. Flies were individually placed in TRI reagent (Ambion) following five days of infection (5 DPI), homogenized total RNA was extracted using Direct-zol RNA miniprep kit, which includes a DNAse step (Zymo Research), reverse-transcribed with M-MLV reverse transcriptase (Promega) and random hexamer primers, and then diluted 1:2 with nuclease-free water. qRT-PCR was performed on an Applied Biosystems StepOnePlus system using Fast SYBR Green Master Mix (Applied Biosystems) and DCV primers, which include 5’-AT rich flaps to improve RT-PCR fluorescent signal (Afonina *et al.*, 2007) (DCV_Forward: 5’

AATAAATCATAAGCCACTGTGATTGATACAACAGAC 3’; DCV_Reverse: AATAAATCATAAGAAGCACGATACTTCTTCCAAACC). We measured the relative fold change in DCV RNA relative to *rp49*, (Dmel_rp49 Forward: 5’ ATGCTAAGCTGTCGCACAAATG 3’; Dmel_rp49 Reverse: 5’ GTTCGATCCGTAACCGATGT

3’). an internal *Drosophila* control gene, calculated as 2^-ΔΔct^ (Livak & Schmittgen, 2001).

### Oral DCV exposure

In separate pilot infections, we determined that a DCV culture diluted to contain approximately 10^5^ DCV RNA copies was enough to establish a viable infection (Figure S1), but did not cause noticeable mortality, and we used this dilution of DCV stock to inoculate all ten DGRP lines. Individual flies were exposed to DCV in vials containing Agar (5% sugar) using 3mL plastic atomizer spray bottles containing 2mL of the sub-lethal DCV dilution. One spray, releasing roughly 50μL of DCV dilution (or sterile Ringer’s solution), was deployed into each vial. Flies were left in the these ‘exposure vials’ for three days to allow them to ingest the viral solution during feeding and grooming, and then tipped into vials containing clean, blue-dyed Lewis medium (see below).

### Survival following infection

Both systemically and orally infected flies were housed individually following infection in vials containing Lewis medium. In the systemic infection experiment, flies were monitored daily for mortality for 38 days post-infection and were transferred to fresh food vials once a week. In the oral infection experiment, flies were transferred to fresh food vials every 3–4 days, and mortality was recorded at this point for the first 32 days post infection and then daily until 40 DPI (oral infection).

### Fecal excretion following oral DCV exposure

Following the exposure period, flies were tipped into vials containing blue-dyed Lewis medium. Blue medium was prepared by adding 0.5g/L FIORI COLORI brilliant blue FCF E133 granules to standard Lewis medium. Flies remained on blue Lewis food for the remainder of the experiment and were tipped to new blue Lewis vials every three to four days. When flies were tipped to new vials, the old vials were kept for fecal spot counts (measured immediately) and fecundity measures (see below). Fecal spots were recorded by photographing vials with a Leica S8APO microscope. A slip of white printer paper (2.5cm × 8.5cm) was inserted into each vial to ensure only spots on one side of the vial were being photographed. These images were then analyzed with ICY image software (Version 1.6.1.1 ICY - Bio Imaging Analysis) and fecal spots were counted using ‘spot detection’ analysis on a 2cm × 4cm region of interest. Each image was checked individually for miscounts, and miscounted spots were removed. Fecal excretion was recorded for 30 days following infection.

### Fecundity

The fecundity of individual flies was measured by counting viable offspring emerging in the vials they were reared in, which happed weekly until day 30 post infection in the systemically infected flies, and every 3–4 days in the orally infected flies, for 28 days following exposure to DCV. Short-term fecundity estimates have been shown to be well correlated with lifetime reproduction in Drosophila (Nguyen & Moehring, 2015) Vials that individuals were tipped from (and following the recording of fecal shedding in the oral infection experiment), were placed in the incubators at 24.5°C ± 0.5 with a 12h:12h light:dark cycle to allow any offspring to develop. After 14 days, the total number of living emerged adult offspring within each vial was recorded as a measure of female fecundity.

### Activity

Locomotor activity was measured using the Drosophila Activity Monitor (DAM2, Trikinetics) as described previously (Pfeiffenberger *et al.*, 2010; Vale & Jardine, 2015). In the DAM, individual fly activity is recorded when individually housed flies break an infrared beam passing through a transparent plastic tube placed symmetrically inside a DAM unit. Activity was measured in a separate experiment on flies reared and exposed to DCV as described above. In systemically infected flies, activity was measured on 10 replicate male flies for each DCV dose the day following septic injury (40 flies in total), and measured for 2 weeks following infection. In the oral infection experiment, activity was recorded for 24 hours, fourteen days after the initial oral exposure. These differences in the timing of activity measurements arise from the faster and more severe effects of systemic infections on locomotor behavior, while we have found that effects on activity following oral infection take longer to manifest, and become apparent 10–15 days after DCV ingestion(Vale & Jardine, 2015). Four replicate flies for each DGRP (10 lines) / sex (M/F) / infection (DCV/Control) combination were tested (160 flies in total). In both experiments, flies were placed individually in a single DAM tube containing a small agar plug on one end, and allocated a slot in one of five DAM unit (each unit can house a maximum of 32 tubes). At least one slot in each DAM unit was filled with an empty tube and at least two slots were left empty as negative controls. All DAM units were placed in the incubator (25 °C 12:12 light:dark cycle) and continuous activity data was collected every minute for 24 hours. Raw activity data was processed using the DAM System File Scan Software (http://www.trikinetics.com) and the resulting data was manipulated using R v. 3.1.3 (The R Foundation for Statistical Computing, Vienna, Austria). Flies that died during the DAM assay (6/40 flies in the systemic infection experiment; 25/160 in the oral infection experiment) were removed from the analysis because they would wrongly bias the estimate of activity.

### Data analysis

All analyses were carried out in JMP 12 (SAS). Survival data was analyzed on the ‘day of death’ using a Cox Proportional Hazards models in with ‘fly sex’ and ‘DCV exposure’ and their interaction as fixed effects (systemic infection experiment) or ‘fly sex’, fly ‘line’ and ‘DCV dose’ and their interactions as fixed effects (oral infection experiment). In the systemic infection, DCV titers were Log_10_- transformed and analyzed in a linear model with ‘DCV Dose’ and ‘Sex’ and their interaction as fixed effects. Fecundity following systemic infection was calculated on the cumulative number of emerged offspring in a model containing ‘DCV dose’ as a fixed effect. In the oral exposure experiment, the cumulative number of offspring was analyzed in a model including ‘Fly line’ and ‘DCV exposure’ and their interaction as fixed effects. Total excretion per fly was analyzed using a linear model with ‘Fly line’, ‘DCV exposure’, and ‘sex’ as categorical fixed effects, ‘Time’ as a continuous covariate, and all pair-wise interactions. Activity was analyzed as the total number of DAM beam breaks recorded per day. Activity following systemic infection was analyzed in a linear model with ‘DCV dose’ and ‘Time’ as fixed effects. Activity following oral infection was measured for 24h and analyzed in a linear model with ‘Fly line’, ‘Sex’ and ‘DCV exposure’ as fixed effects. In all analyses, individual replicate was included as a random factor, and in all cases accounted for only 2–5% of the total variance.

## RESULTS

### Experiment 1 Sub-lethal systemic infection

In a first experiment, we tested how systemic infection with very low sub-lethal doses of DCV affected fly health. The survival of both female and male flies exposed to doses of 10^2^ and 10^3^ DCV IU/ ml did not differ from control flies that had been pricked with sterile buffer solution (Figure 1a). In females, 100% flies exposed to these doses survived infection during the 38-day survival assay, while roughly 20% of males died during this period (Figure 1a. However, this difference in survival between sexes (‘sex’ effect, Table 1), was also observed in control flies and therefore is likely to reflect sex-specific responses to injury during intra-thoracic pricking than to infection. Flies infected with a slightly higher concentration of 10^5^ DCV IU/ ml died significantly faster than control flies. This virus concentration-specific pattern of mortality was generally consistent with the observed DCV titers measured 5 days following infection, (Table 2, ‘dose’ effect) which were generally higher in male flies across all DCV concentrations (Table 2, ‘sex’ effect, Figure 1b). Our experiment therefore spanned the range of sub-lethal viral doses, with 10^5^ DCV IU/ ml being the lowest virus concentration with lethality in the experiment (Figure 1a).

**Table 1.**
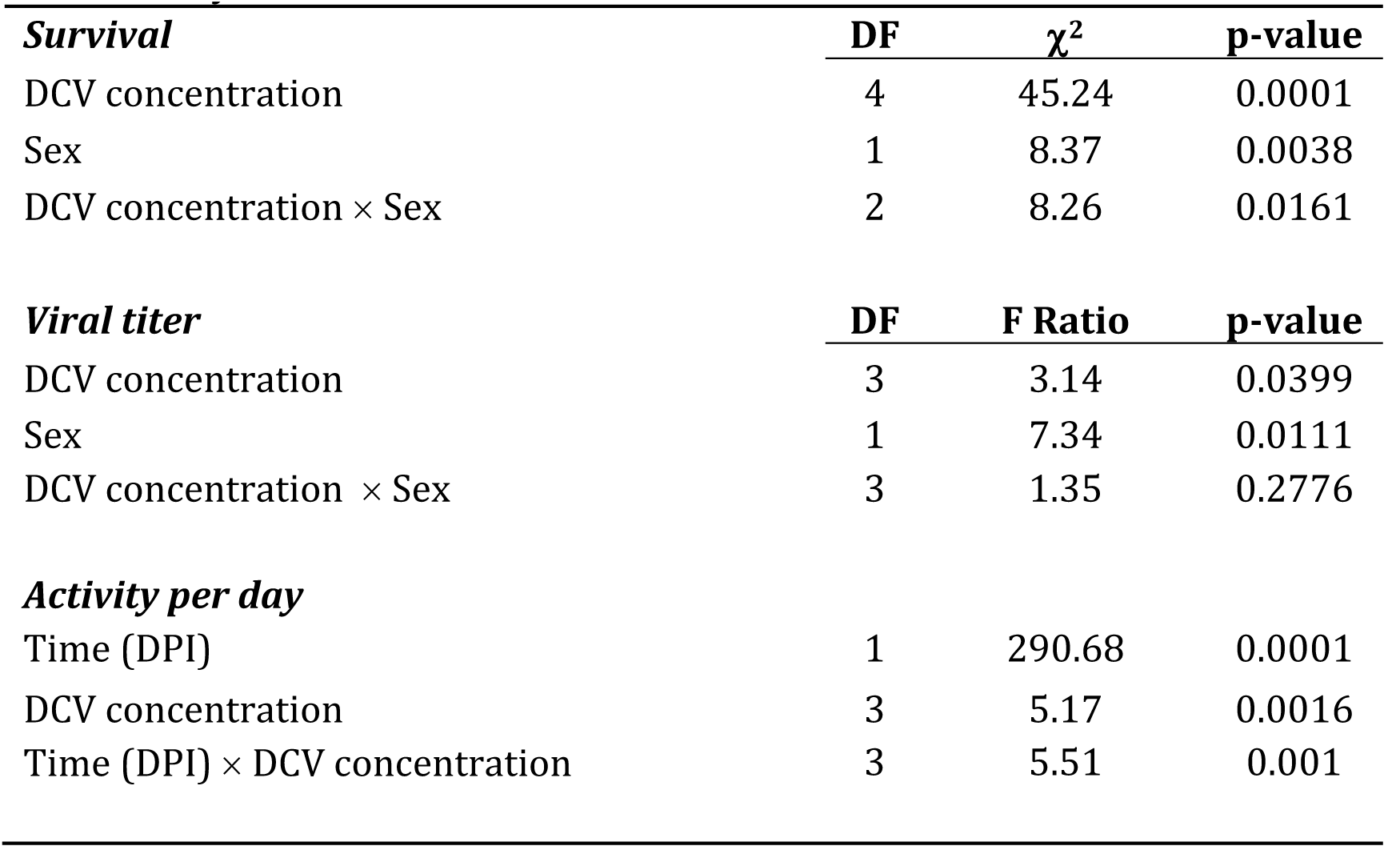
Systemic infection

**Table 2.** Oral infection

|  | **DF** | **F Ratio** | **p-value** |
| --- | --- | --- | --- |
| ***Fecundity*** |  |  |  |
| DGRP Line | 9 | 16.17 | <.0001 |
| DCV exposure | 1 | 0.99 | 0.3186 |
| DGRP Line × DCV exposure | 9 | 2.59 | 0.0076 |
| ***Activity per day*** |  |  |  |
| DGRP Line | 9 | 2.91 | 0.0037 |
| Sex | 1 | 0.02 | 0.8947 |
| DCV exposure | 1 | 1.45 | 0.2315 |
| DGRP Line × Sex | 9 | 2.18 | 0.0277 |
| DGRP Line × DCV exposure | 9 | 0.67 | 0.7352 |
| Sex × DCV exposure | 1 | 0.12 | 0.7244 |
| ***Fecal excretion*** |  |  |  |
| DGRP Line | 9 | 32.17 | 0.0001 |
| Sex | 1 | 212.66 | 0.0001 |
| Time (DPI) | 1 | 29.95 | 0.0001 |
| DCV exposure | 1 | 72.83 | 0.0001 |
| DGRP Line × DCV exposure | 9 | 4.46 | 0.0001 |
| Sex × DCV exposure | 1 | 13.45 | 0.0003 |
| Time (DPI) × DCV exposure | 1 | 0.23 | 0.6295 |
| DGRP Line × Sex | 9 | 31.22 | 0.0001 |
| DGRP Line × Time (DPI) | 9 | 1.28 | 0.2405 |
| Sex × Time (DPI) | 1 | 0.06 | 0.806 |

**Figure 1.**
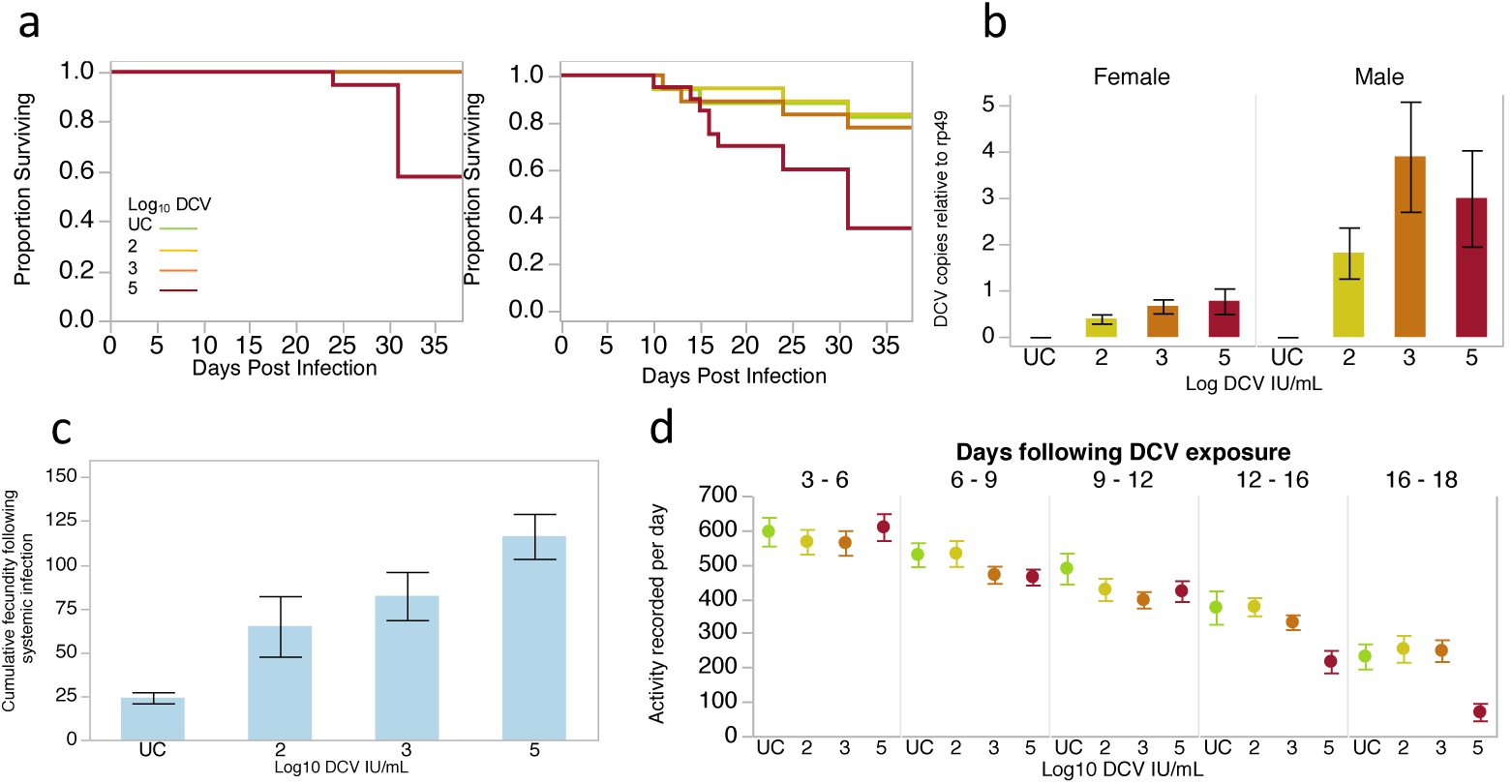
Sub-lethal systemic infection, 1a. Kaplan-Meier curves showing the survival of 20 replicate flies exposed systemically to sub-lethal concentrations of DCV. 1b. DCV titers measured in male and female flies relative to an internal control gene (rp49), following 3 days of systemic infection with sub-lethal concentrations of DCV. For each DCV concentration, data are the average of duplicate qPCR reactions for 5 individual flies, lc. Total fecundity recorded for 30 days following systemic infection in mated females. Data are the means ± SE of 18–19 replicate females flies. 1d. Daily locomotor activity of male flies following systemic infection with DCV. Data are 3 day averages of 7–10 replicate flies for each inoculation concentration. UC are uninfecred controls.

#### Fecundity following systemic DCV infection

We used mated females, which allowed us to quantify fly reproductive health during systemic infection by following the number of adult offspring produced by individual females for 30 days following infection. The total fecundity measured during this period varied according to the dose females had received (F3,66 = 10.32, p<0.0001) and we observed that the total reproduction of infected flies was higher than control flies, and increased in a dose-specific manner (Figure 1c).

#### Activity following systemic DCV infection

The locomotor activity of individual male flies infected systemically with all sub-lethal concentrations of DCV was measured during 18 days after infection in a Trikinetics^®^ Drosophila Activity Monitor (DAM). All flies included in the analysis remained alive for the whole period, so changes in activity were not confounded with potential death of individual flies. We found that flies in all treatments, including uninfected controls, showed a reduction in activity over the course of the activity assay (Figure 1d, Table time effect). This general effect is not especially surprising given the constrained environment experienced by flies in the DAM tubes, and that the only source of nutrition and hydration is small agar plug. However, our analysis showed that the temporal reduction in activity depended on the dose that flies had received (‘time x dose’ interaction, Table 1). In the early stages of infection flies receiving the higher of the 4 doses (10^3^ and 10^5^ DCV copies) showed a reduction in activity relative to control flies and those receiving the lowest dose. Over time, a reduction in locomotor activity was most apparent in flies infected with the highest dose of 10^5^ DCV copies (Figure 1d).

### Experiment 2 Sub-lethal gut infection

In a separate experiment, we tested how exposure to a single sub-lethal dose of DCV through the oral route of infection impacted upon fly health. We conducted the experiment on ten fly lines from the DGRP panel (Mackay *et al.*, 2012) and we included both male and female flies to test for the effects of host genetic background and sex in response to sub-lethal oral infection. While DGRP lines differ in their lifespan in the absence of infection (Durham *et al.*, 2014), we did not detect any difference between DGRP lines or between sexes in their survival during oral DCV infection compared to control flies (Table S1) which, as expected, was generally non-lethal across all lines.

#### Fecundity following oral exposure to DCV

Despite not observing any effects on fly survival during infection, we detected significant variation in reproductive health following exposure to DCV. The total fecundity of females during the 28 days following oral exposure to DCV (or a control inoculum) varied significantly between DGRP lines (Figure 2; Table 2), reflecting well-known genetic differences in the lifetime reproductive output of these lines (Durham *et al.*, 2014). In addition, we found line-specific fecundity responses to DCV infection (‘infection status x line’, Table 2, see also Table S2 for pairwise contrasts). In some lines (158, 491, 317) low-level oral infection resulted in a decrease in fecundity; in other lines (821, 358) there was no detectable effect of DCV exposure; while in 2 lines we detected significant increases in fecundity in DCV infected flies compared to uninfected control flies of the same genetic background (Figure 2; see Table S2 for least-square pairwise contrasts).

**Figure 2.**
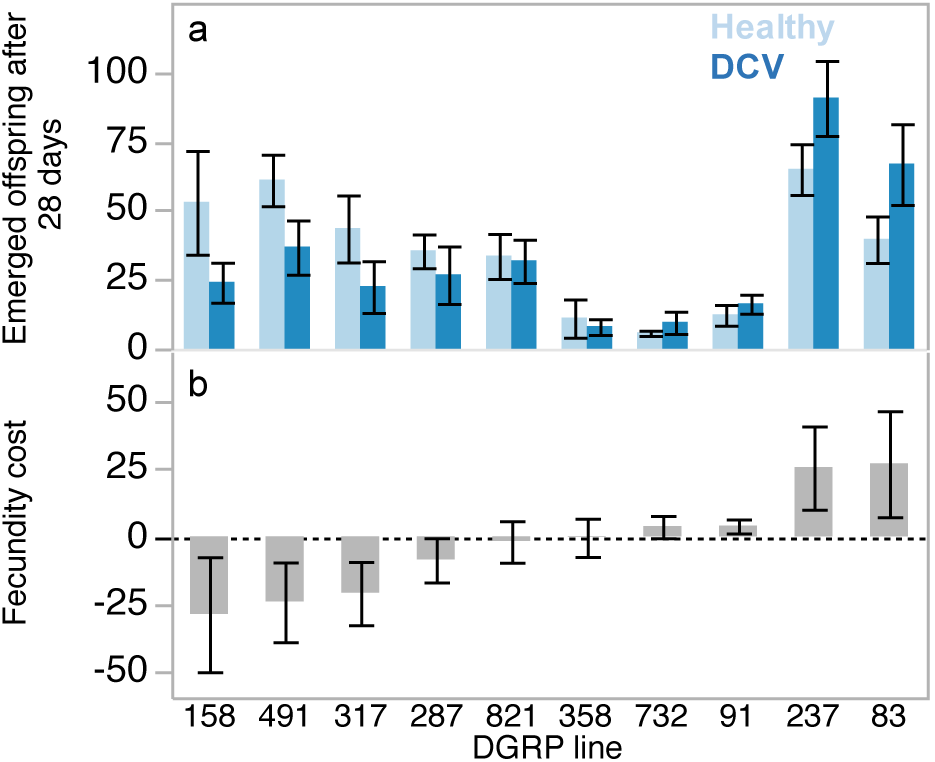
Fecundity following oral DCV exposure. 2a. The cumulative number of adult offspring from healthy (light bars) or DCV-exposed (dark bars) single female flies over the course of the 28-day experiment. 2b. Shows the fecundity difference between healthy and infected flies for the same 10 DRGP lines. In both plots, DGRP lines are ordered from the greatest decrease to the highest fecundity increase. Significant pairwise contrasts (reported in Table S2) are indicated by asterisks. Data are the mean ± SE of eight individual replicate females.

#### Locomotor activity following oral exposure to DCV

Overall, DGRP lines differed in their activity in a sex specific way (‘Fly line x Sex’ effect Table 2), but these differences were not altered by infection. While we detected a reduction in locomotor activity following systemic infection (Figure 1d), we did not detect any effect of oral DCV exposure on the overall activity of flies (Table 2, Figure 3).

**Figure 3.**
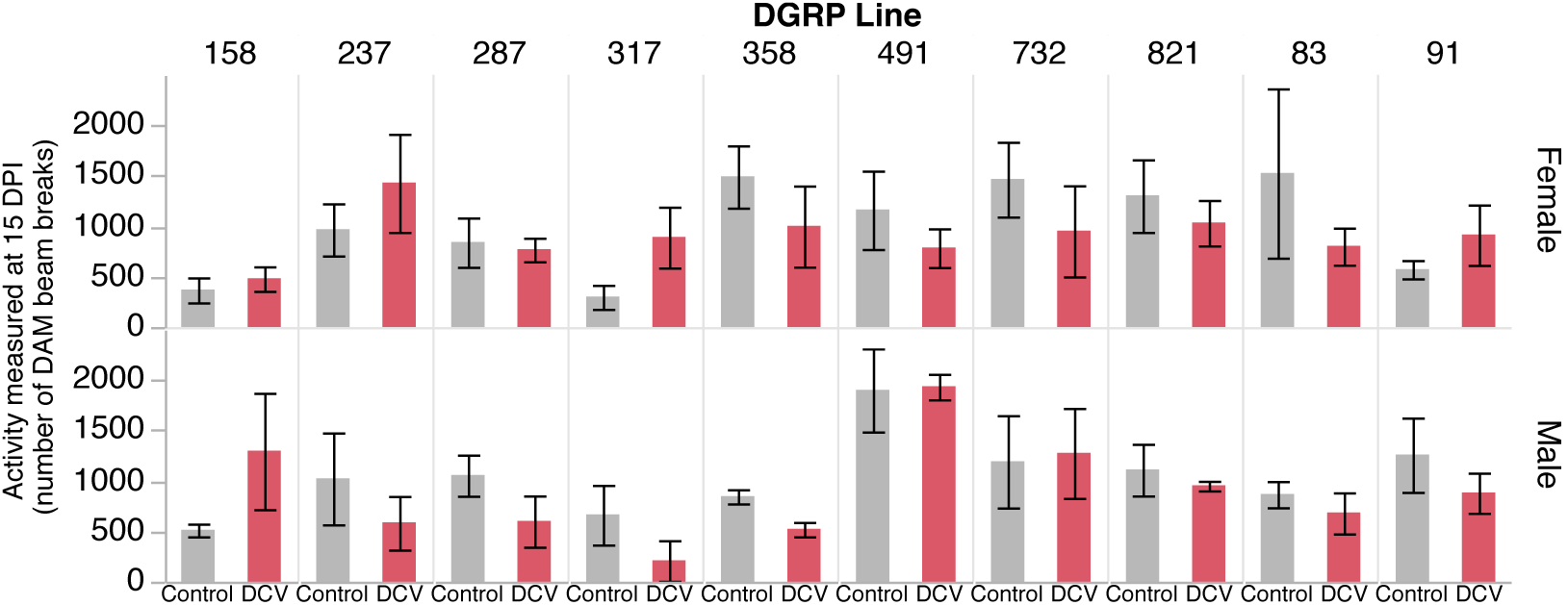
Locomotor activity following oral DCV exposure. Data show mean ± SE activity of four replicate flies per sex and DGRP line, measured for 24 hours 14 days following exposure to DCV (red) or uninfecred controls (grey).

#### Fecal excretion following oral exposure to DCV

We quantified fecal excretion for 30 days following DCV exposure as a proxy for gut health, by counting fecal spots excreted into vials after ingestion of blue-dyed food. Overall we found that males showed higher levels of fecal excretion compared to females (Table 2, ‘sex’ effect; Figure 4) and that DCV infection was associated with a general reduction in fecal excretion throughout the 30-day observation period (‘Infection status’ effect, Figure 4). However, we found that males and females differed in the overall severity of this reduction (‘sex x infection status’ effect), with males showing a greater reduction in defecation overall (Figure 4). Furthermore, we found significant variation among the DGRP lines in the magnitude of the effect of DCV on fecal excretion (‘fly line x infection status’ effect).

**Figure 4.**
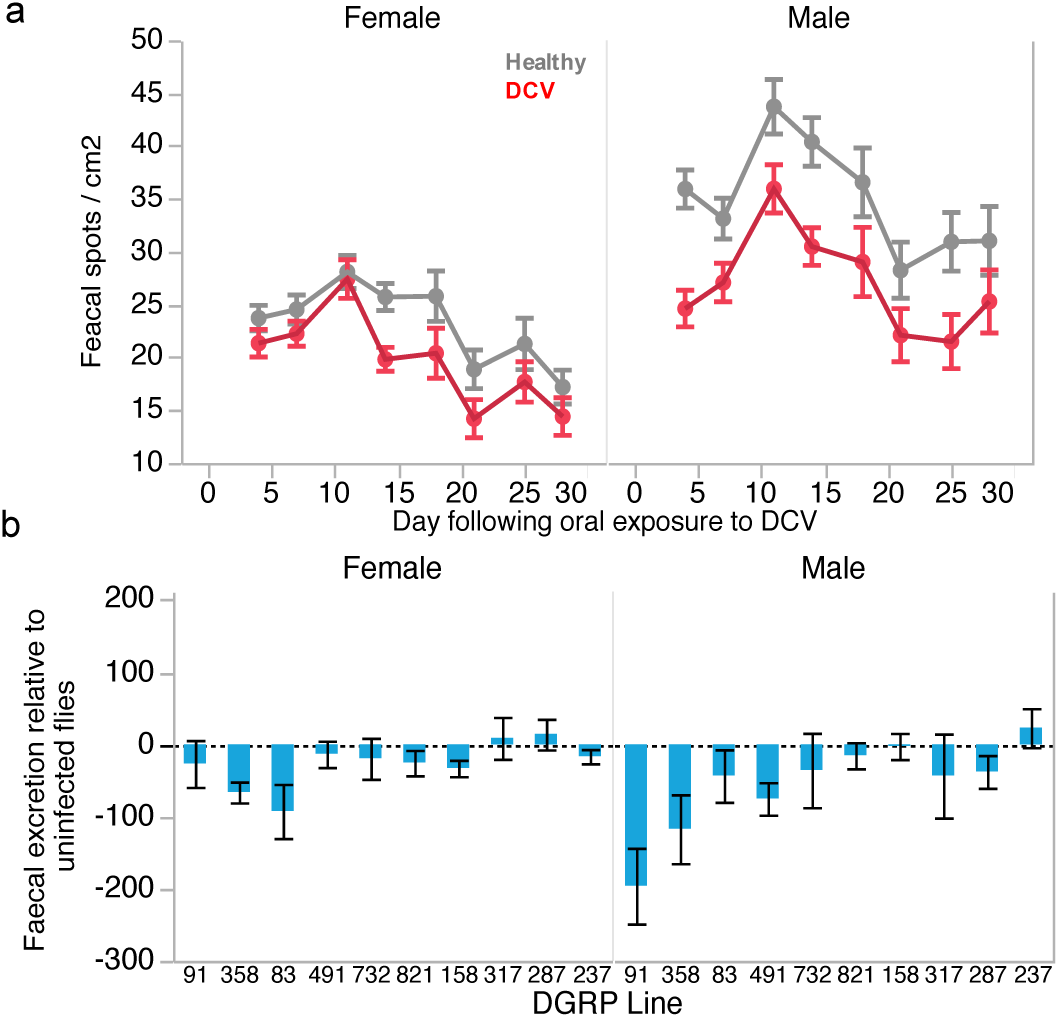
Fecal excretion following oral DCV exposure. 4a. The general effect of DCV exposure (red) or a control inoculum (grey) on the number of fecal spots shed over time. Data are plotted separately for males and females. Each time point is the mean ± SE of 8 replicate individual flies averaged across all 10 DGRP lines. 4b. Shows the difference between control and infected flies for each DRGP line. Data are the mean ± SE of eight individual replicate flies for each sex and line combination.

## DISCUSSION

We find that sub-lethal infections with DCV can cause measurable morbidity that is relevant for the fitness costs experienced by *D. melanogaster* during DCV infection. In systemic and oral infections with doses that did not cause mortality we observed DCV-induced effects on fly reproductive output, digestive health, and locomotor activity.

### Systemically infected flies increase reproductive output

We found that the fly line used in the systemic infection experiment showed an increase in reproductive output when infected with sub-lethal doses of DCV. There are numerous examples from both invertebrates and vertebrates of fecundity increases following infection (Bonneaud *et al.*, 2004; Vale & Little, 2012; Leventhal *et al.*, 2014;Vézilier *et al.*, 2015). In addition, earlier work reported that DCV infection could increase ovariole number and decrease development time in *D. melanogaster* (Thomas-Orillard, 1984; Gomariz-Zilber & Thomas-Orillard, 1993). However, a subsequent re-analysis of these data showed very weak support for the beneficial effects of DCV infection (Longdon, 2015). It is notable however that neither of the earlier studies measured the number of viable offspring of infected flies compared to healthy ones. The fecundity data we report therefore suggests that DCV may indeed result in increased reproductive output.

A dose-dependent increase in fecundity could suggest a direct effect of DCV infecting fly ovaries, but it is unclear why such a strategy would be adaptive for the virus. An alternative hypothesis may instead involve more complex interactions between the allocation of resources during DCV infection, and how they relate to fly nutritional stress and reproductive investment. For example, *D. melanogaster* females selected under conditions of nutritional stress were found to produce a greater number of ovarioles, while the offspring of starved mothers also exhibited greater investment in reproduction (Wayne *et al.*, 2006). Similar to the studies cited above, this work also focused on ovariole number and egg production, and did not quantify female lifetime fecundity. Given that DCV infection is known to lead to intestinal obstruction, one possibility for the increase in the number of adult offspring we observed in infected flies is that DCV-induced nutritional stress leads to a greater production of ovarioles, and consequently, an increased number of offspring. Given we only tested a single fly line however, it important to note that this response may not be universal. As we discuss below fecundity responses to infection have generally been found to differ between host genotypes (Vale & Little, 2012; Parker *et al.*, 2014)

### Fecundity costs and benefits of DCV infection are genotype-specific

Similar to systemically infected flies (Figure 1c), we also find evidence for fecundity benefits in orally exposed flies, but these benefits were only revealed in two out of the ten genetic backgrounds we tested. Indeed, in three of the tested lines, DCV infection resulted in lower reproductive output. Taking fecundity as a proxy for evolutionary fitness, the existence of genotype specific fitness costs and benefits means that DCV could be a potentially powerful driver of *D. melanogaster* evolutionary dynamics. Previous analyses of *Drosophila* spp. population genetic data have shown that the fastest evolving *D. melanogaster* genes are those involved in RNAi-based antiviral defense (Obbard *et al.*, 2006, 2009; Early *et al.*, 2016), but the DCV-induced fitness costs that drive this rapid evolution in wild-infected flies (where infections are persistent and often nonlethal), has remained obscure. These data suggest that genotype-specific fecundity costs and benefits of DCV infection could potentially mediate the arms-race between flies and viruses.

### Systemically infected flies show a dose-dependent decline in activity over time

Reduced activity, or lethargy, following infection is a common response to infection across a range of taxa (Hart, 1988; Adelman & Martin, 2009; Sullivan *et al.*, 2016). The most obvious explanation for reduced activity is simply that infected individuals are sick, and lethargy reflects the underlying pathology of infection (Moore, 2013). A popular alternative explanation is that infection-induced lethargy evolved as an adaptive host strategy that conserves energy, which may then be allocated to other physiological tasks such as mounting an immune response (Hart, 1988; Adelman & Martin, 2009).

Support for the adaptive nature of these ‘sickness behaviours’ has come mainly from vertebrate species challenged with deactivated pathogens or their derived components, which are sufficient to stimulate an immune response without causing pathology (Adelman & Martin, 2009; Lopes *et al.*, 2016). In addition to vertebrates, sickness behaviors including lethargy and anorexia have also been described in insect hosts (Ayres & Schneider, 2009; Kazlauskas *et al.*, 2016;Sullivan *et al.*, 2016). However, in the current experiment it is not possible to disentangle the effect of an adaptive sickness behavior from the direct effect of pathology caused by replicating DCV. Regardless of the underlying cause of reduced activity, it is likely to come at an additional cost of lower involvement in fitness-enhancing activities such as foraging, competing for resources with conspecifics, or courtship and mating (Adelman & Martin, 2009; Adamo *et al.*, 2015; Vale & Jardine, 2016). Further, reduced activity following infection can also reduce the potential for disease spread (Lopes *et al.*, 2016). In the context of understanding sub-lethal DCV infection in an ecological setting, reduced activity may therefore be a potentially important source of DCV-induced fitness costs and benefits.

We did not find an effect of oral DCV exposure on fly activity. Previous work has shown that *Drosophila*, especially females, show a reduction in activity following oral infection with DCV (Vale & Jardine, 2015). However, the viral concentration that flies were exposed to in that experiment was at least 1000x higher, so it is likely that in the current experiment flies did not ingest virus in quantities large enough to affect locomotor activity.

### The severity of DCV-induced digestive dysfunction is sex-specific

Previous work has shown that DCV infection results in digestive dysfunction, leading to increased body mass due to the inability to excrete digested food (Arnold *et al.*, 2013; Chtarbanova *et al.*, 2014). We found that this measure of gut health varied between genotypes and also between sexes. Extensive genetic variation for gut immune-competence has previously been reported in the DGRP panel (Bou Sleiman *et al.*, 2015), which could underlie some of the variation we observe in DCV-associated digestive dysfunction in some lines. Although that study focused on enteric infection with entomopathogenic bacteria, the mechanisms that mediate variation in gut health during infection include general processes of gut damage and repair, such as the production of reactive oxygen species (ROS) and the production of intestinal stem cells during epithelial repair (Buchon *et al.*, 2013). It is plausible that these mechanisms also mediate disease severity during enteric virus infection, but we are unaware of any systematic study of genetic variation in gut immune-competence during viral infection.

### Concluding remarks

Altogether, these measures of sub-lethal morbidity give insight into the potential fitness costs of low-level, persistent DCV infection in Drosophila. More generally, the combination of both positive and negative effects on fly fitness effects according to the specific host genetic background presents a non-trivial evolutionary scenario for host immune defense (Gandon & Vale, 2014). For instance, frequent encounters between beneficial symbionts and detrimental pathogens are hypothesized to have played a role in the evolution of aphid immune systems, which lack several components of the IMD immune pathway critical for the recognition and elimination of Gram-negative bacteria (Gerardo et al., 2010). The combination of fitness costs and benefits of infection, such as those incurred during DCV infection, may therefore have driven the evolution of immune defense across a wide range of host taxa, from insects to mammals (Elsik, 2010; Gerardo et al., 2010; Lee & Mazmanian, 2010; Gandon & Vale, 2014).

## Acknowledgements

We are grateful to D. Obbard and lab members (Edinburgh) for general technical support and for advice about DCV infection. We also thank H. Cowan and H. Borthwick for help with media preparation. This work was supported by a strategic award from the Wellcome Trust for the Centre for Immunity, Infection and Evolution (http://ciie.bio.ed.ac.uk; grant reference no. 095831), and by a Society in Science - Branco Weiss fellowship (http://www.society-in-science.org), both awarded to P. Vale. All authors declare no conflict of interest.

## Supplementary File for **Costs and benefits of sub-lethal Drosophila C Virus infection**

This file contains:

– Table S1. Cox proportional hazards analysis of survival following oral exposure to DCV.
– Table S2. Least Square Means Student’s t pairwise contrasts between exposed and control fecundity following oral DCV exposure.
– Figure S1. DCV increases in titer following oral exposure to approximately 10^5^ DCV copies.

**Table S1.** Output of Cox proportional hazard model testing variation in survival following oral infection.

| Survival during oral infection | DF | $\chi^2$ | p-value |
| --- | --- | --- | --- |
| DGRP Line | 9 | 3.87084122 | 0.9197 |
| Sex | 1 | 3.82E-07 | 0.9995 |
| DGRP Line*Sex | 9 | 2.85864198 | 0.9696 |
| Infection status | 1 | 4.73E-08 | 0.9998 |
| DGRP Line* Infection status | 9 | 0.74383375 | 0.9998 |
| Sex* Infection status | 1 | 1.07E-06 | 0.9992 |
| DGRP Line*Sex* Infection status | 9 | 0.25051421 | 1 |

**Table S2.**
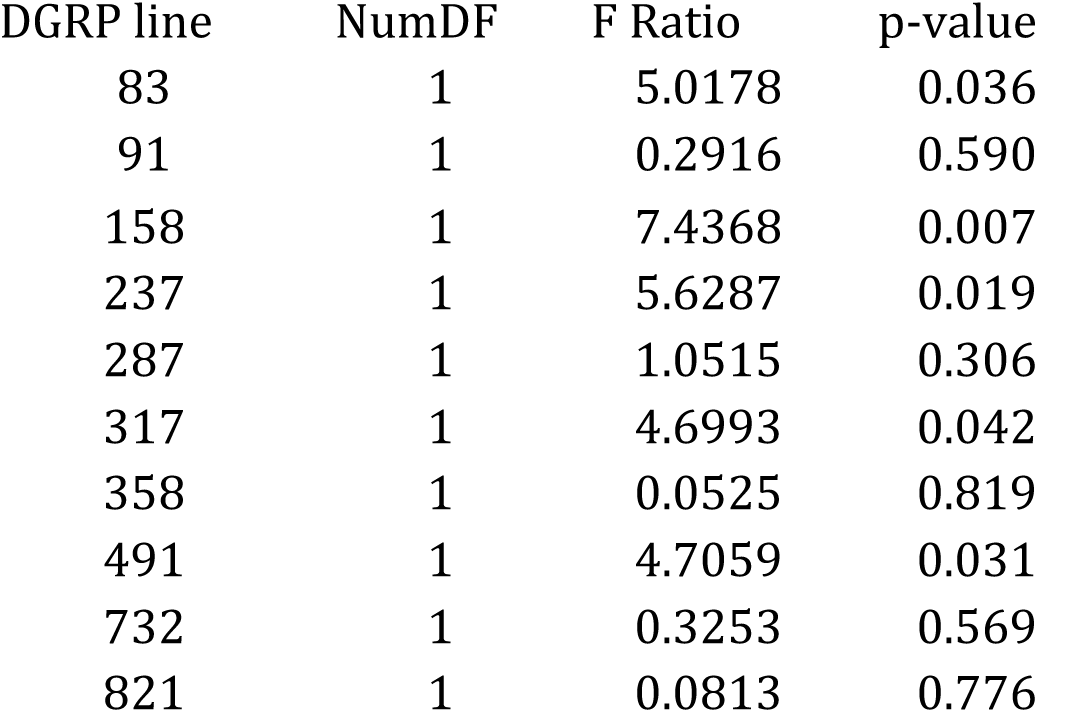
Least Square Means Student’s t pairwise contrasts between exposed and control fecundity following oral DCV exposure

| DGRP line | NumDF | F Ratio | p-value |
| --- | --- | --- | --- |
| 83 | 1 | 5.0178 | 0.036 |
| 91 | 1 | 0.2916 | 0.590 |
| 158 | 1 | 7.4368 | 0.007 |
| 237 | 1 | 5.6287 | 0.019 |
| 287 | 1 | 1.0515 | 0.306 |
| 317 | 1 | 4.6993 | 0.042 |
| 358 | 1 | 0.0525 | 0.819 |
| 491 | 1 | 4.7059 | 0.031 |
| 732 | 1 | 0.3253 | 0.569 |
| 821 | 1 | 0.0813 | 0.776 |

**Figure S1.**
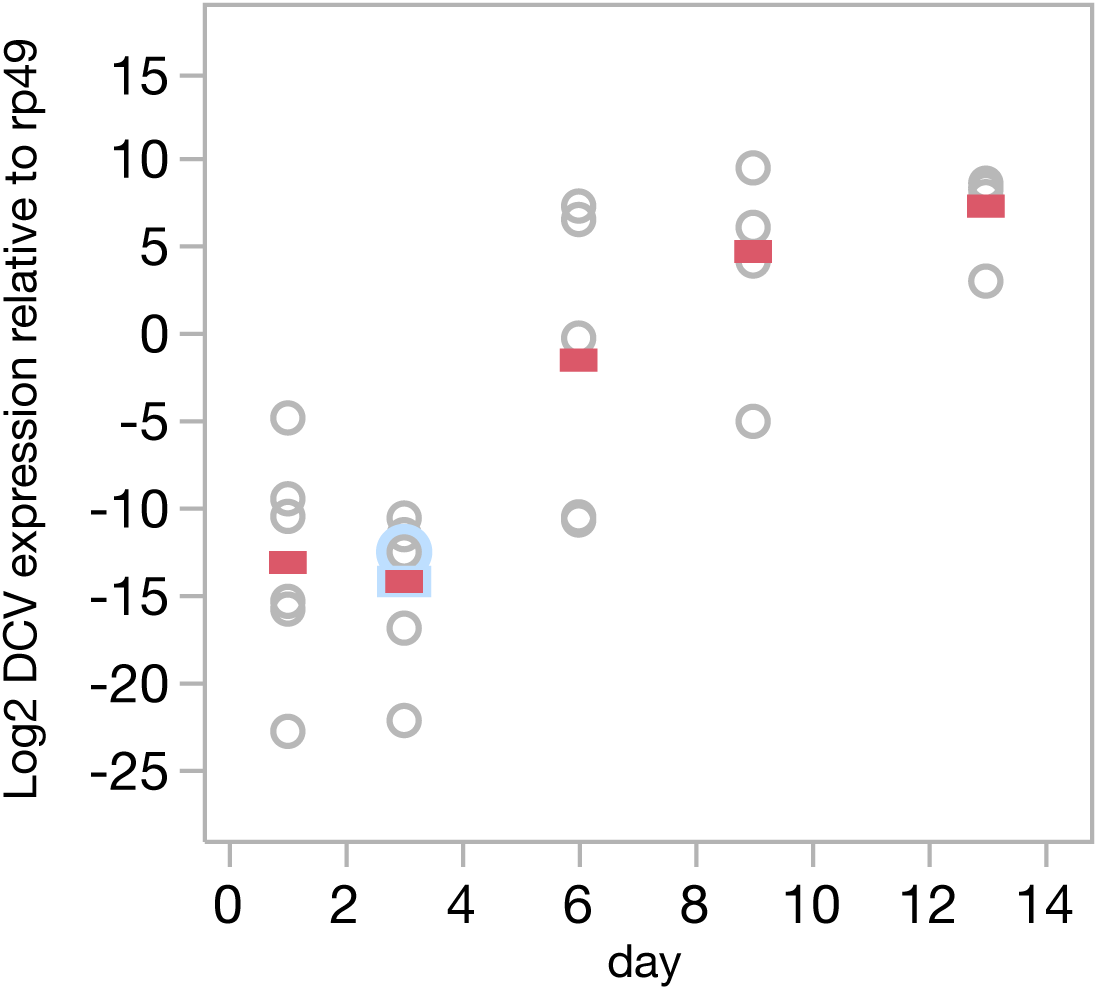
DCV increases in titer following oral exposure to with approximately 10^5^ DCV copies (F1,27 = 57.97, p< 0.001). This experiment was carried out in *D. melanogaster* OreR. Data show the Log2 DCV expression relative to an internal Drosophila control gene (rp49), measured in six individual female flies at each time point following exposure. Oral exposure to DCV was carried out as described in the main text.

## REFERENCES

Adamo, S.A., Gomez-Juliano, A., LeDue, E.E., Little, S.N. & Sullivan, K. 2015. Effect of immune challenge on aggressive behaviour: how to fight two battles at once. Anim. Behav. 105: 153–61.

Adelman, J.S. & Martin, L.B. 2009. Vertebrate sickness behaviors: Adaptive and integrated neuroendocrine immune responses. Integr. Comp. Biol. 49: 202–214.

Afonina, I., Ankoudinova, I., Mills, A., Lokhov, S., Huynh, P. & Mahoney, W. 2007. Primers with 5’ flaps improve real-time PCR. BioTechniques 43: 770, 772, 774.

Ambrose, R.L., Lander, G.C., Maaty, W.S., Bothner, B., Johnson, J.E. & Johnson, K.N. 2009. Drosophila A virus is an unusual RNA virus with a T=3 icosahedral core and permuted RNA-dependent RNA polymerase. J. Gen. Virol. 90: 2191–2200.

Arnold, P.A., Johnson, K.N. & White, C.R. 2013. Physiological and metabolic consequences of viral infection in Drosophila melanogaster. J. Exp. Biol. 216: 3350–3357.

Ayres, J.S. & Schneider, D.S. 2009. The Role of Anorexia in Resistance and Tolerance to Infections in Drosophila. PLoS Biol. 7: e1000150.

Bonneaud, C., Mazuc, J., Chastel, O., Westerdahl, H. & Sorci, G. 2004. Terminal investment induced by immune challenge and fitness traits associated with major histocompatibility complex in the house sparrow. Evol. Int. J. Org.Evol. 58: 2823–2830.

Bou Sleiman, M.S., Osman, D., Massouras, A., Hoffmann, A.A., Lemaitre, B. & Deplancke, B. 2015. Genetic, molecular and physiological basis of variation in Drosophila gut immunocompetence. Nat. Commun. 6.

Buchon, N., Broderick, N.A. & Lemaitre, B. 2013. Gut homeostasis in a microbial world: insights from Drosophila melanogaster. Nat. Rev. Microbiol. 11: 615–626.

Chtarbanova, S., Lamiable, O., Lee, K.-Z., Galiana, D., Troxler, L., Meignin, C., et al. 2014. Drosophila C virus systemic infection leads to intestinal obstruction. J. Virol., doi: 10.1128/JVI.02320–14.

Conway, M.J., Colpitts, T.M. & Fikrig, E. 2014. Role of the Vector in Arbovirus Transmission. Annu. Rev. Virol. 1: 71–88.

Dostert, C., Jouanguy, E., Irving, P., Troxler, L., Galiana-Arnoux, D., Hetru, C., et al. 2005. The Jak-STAT signaling pathway is required but not sufficient for the antiviral response of drosophila. Nat. Immunol. 6: 946–953.

Dow, J.A.T. & Davies, S.A. 2001. The Drosophila melanogaster malpighian tubule. In: (B. -A. in I. Physiology, ed), pp. 1–83. Academic Press.

Durham, M.F., Magwire, M.M., Stone, E.A. & Leips, J. 2014. Genome-wide analysis in Drosophila reveals age-specific effects of SNPs on fitness traits. Nat. Commun. 5.

Dwyer, G., Dushoff, J. & Yee, S.H. 2004. The combined effects of pathogens and predators on insect outbreaks. Nature 430: 341–345.

Early, A.M., Arguello, J.R., Cardoso-Moreira, M., Gottipati, S., Grenier, J.K. & Clark, A.G. 2016. Survey of Global Genetic Diversity Within the Drosophila Immune System. Genetics, doi: 10.1534/genetics.116.195016.

Elsik, C.G. 2010. The pea aphid genome sequence brings theories of insect defense into question. Genome Biol. 11: 106.

Ferreira, Á.G., Naylor, H., Esteves, S.S., Pais, I.S., Martins, N.E. & Teixeira, L. 2014. The Toll-Dorsal Pathway Is Required for Resistance to Viral Oral Infection in Drosophila. PLoS Pathog. 10.

Gandon, S. & Vale, P.F. 2014. The evolution of resistance against good and bad infections. J. Evol. Biol. 27: 303–312.

Gerardo, N.M., Altincicek, B., Anselme, C., Atamian, H., Barribeau, S.M., Vos, M. de, et al. 2010. Immunity and other defenses in pea aphids, Acyrthosiphon pisum. Genome Biol. 11: R21.

Gomariz-Zilber, E., Poras, M. & Thomas-Orillard, M. 1995. Drosophila C virus: experimental study of infectious yields and underlying pathology in Drosophila melanogaster laboratory populations. J. Invertebr. Pathol. 65: 243–247.

Gomariz-Zilber, E. & Thomas-Orillard, M. 1993. Drosophila C virus and Drosophila hosts: a good association in various environments. J. Evol. Biol. 6: 677–689.

Habayeb, M.S., Cantera, R., Casanova, G., Ekström, J.-O., Albright, S. & Hultmark, D. 2009. The Drosophila Nora virus is an enteric virus, transmitted via feces. J. Invertebr. Pathol. 101: 29–33.

Hart, B.L. 1988. Biological basis of the behavior of sick animals. Neurosci. Biobehav. Rev. 12: 123–137.

Huszar, T. & Imler, J. 2008. Drosophila Viruses and the Study of Antiviral Host-Defense. In: Advances in Virus Research, pp. 227–265. Academic Press.

Huylmans, A.K. & Parsch, J. 2014. Population- and Sex-Biased Gene Expression in the Excretion Organs of Drosophila melanogaster. G3 GenesGenomesGenetics 4:2307–2315.

Jousset, F.-X., Bergoin, M. & Revet, B. 1977. Characterization of the Drosophila C Virus. J. Gen. Virol. 34: 269–283.

Kapun, M., Nolte, V., Flatt, T. & Schlötterer, C. 2010. Host Range and Specificity of the Drosophila C Virus. PLoS ONE 5: e12421.

Karlikow, M., Goic, B. & Saleh, M.-C. 2014. RNAi and antiviral defense in Drosophila: Setting up a systemic immune response. Dev. Comp. Immunol. 42: 85–92.

Kazlauskas, N., Klappenbach, M., Depino, A.M. & Locatelli, F.F. 2016. Sickness Behavior in Honey Bees. Front. Physiol. 7.

Kemp, C. & Imler, J.-L. 2009. Antiviral immunity in drosophila. Curr. Opin. Immunol. 21: 3–9.

Lacey, L.A., Grzywacz, D., Shapiro-Ilan, D.I., Frutos, R., Brownbridge, M. & Goettel, M.S. 2015. Insect pathogens as biological control agents: Back to the future. J. Invertebr. Pathol. 132: 1–41.

Lautié-Harivel, N. & Thomas-Orillard, M. 1990. Location of Drosophila C virus target organs in Drosophila host population by an immunofluorescence technique. Biol. Cell 69: 35–39.

Lee, Y.K. & Mazmanian, S.K. 2010. Has the Microbiota Played a Critical Role in the Evolution of the Adaptive Immune System? Science 330: 1768–1773.

Leventhal, G.E., Dünner, R.P. & Barribeau, S.M. 2014. Delayed virulence and limited costs promote fecundity compensation upon infection. Am. Nat. 183: 480–493.

Lewis, E. 2014. A new standard food medium. 1960 Drosophila Information Service. in. Cold Spring Harb. Protoc. 2014: pdb.rec081414.

Livak, K.J. & Schmittgen, T.D. 2001. Analysis of relative gene expression data using real-time quantitative PCR and the 2(-Delta Delta C(T)) Method. Methods San Diego Calif 25: 402–408.

Longdon, B. 2015. Examination of data claiming Drosophila C virus is beneficial do not support this claim. *Figshare*, doi:http://dx.doi.org/10.6084/m9.figshare.1297064.

Longdon, B., Cao, C., Martinez, J. & Jiggins, F.M. 2013. Previous Exposure to an RNA Virus Does Not Protect against Subsequent Infection in Drosophila melanogaster. PLoS ONE 8: e73833.

Lopes, P.C., Block, P. & König, B. 2016. Infection-induced behavioural changes reduce connectivity and the potential for disease spread in wild mice contact networks. Sci. Rep. 6: 31790.

Mackay, T.F.C., Richards, S., Stone, E.A., Barbadilla, A., Ayroles, J.F., Zhu, D., et al. 2012. The Drosophila melanogaster Genetic Reference Panel. Nature 482: 173–178.

Magwire, M.M., Fabian, D.K., Schweyen, H., Cao, C., Longdon, B., Bayer, F., et al. 2012. Genome-Wide Association Studies Reveal a Simple Genetic Basis of Resistance to Naturally Coevolving Viruses in Drosophila melanogaster. PLoS Genet 8: e1003057.

Merkling, S.H., Bronkhorst, A.W., Kramer, J.M., Overheul, G.J., Schenck, A. & Van Rij, R.P. 2015. The Epigenetic Regulator G9a Mediates Tolerance to RNA Virus Infection in Drosophila. PLoS Pathog 11: e1004692.

Miller, L.K. & Ball, L.A. (eds). 1998. The Insect Viruses. Springer US, Boston, MA.

Moore, J. 2013. An overview of parasite-induced behavioral alterations – and some lessons from bats. J. Exp. Biol. 216: 11–17.

Nayak, A., Tassetto, M., Kunitomi, M. & Andino, R. 2013. RNA Interference-Mediated Intrinsic Antiviral Immunity in Invertebrates. In: *Intrinsic Immunity* (B. R. Cullen, ed), pp. 183–200. Springer Berlin Heidelberg.

Nguyen, T.T.X. & Moehring, A.J. 2015. Accurate Alternative Measurements for Female Lifetime Reproductive Success in Drosophila melanogaster. PLOS ONE 10: e0116679.

Obbard, D.J., Jiggins, F.M., Halligan, D.L. & Little, T.J. 2006. Natural selection drives extremely rapid evolution in antiviral RNAi genes. Curr. Biol. CB 16: 580–585.

Obbard, D.J., Welch, J.J., Kim, K.-W. & Jiggins, F.M. 2009. Quantifying Adaptive Evolution in the Drosophila Immune System. PLoS Genet. 5: e1000698.

Parker, B.J., Garcia, J.R. & Gerardo, N.M. 2014. Genetic variation in resistance and fecundity tolerance in a natural host-pathogen interaction. Evol. Int. J. Org. Evol. 68: 2421–2429.

Pfeiffenberger, C., Lear, B.C., Keegan, K.P. & Allada, R. 2010. Locomotor Activity Level Monitoring Using the Drosophila Activity Monitoring (DAM) System. Cold Spring Harb. Protoc. 2010: pdb.prot5518.

Reed, L.J. & Muench, H. 1938. A Simple Method of Estimating Fifty Per Cent Endpoints,. Am. J. Epidemiol. 27: 493–497.

Rosario, K. & Breitbart, M. 2011. Exploring the viral world through metagenomics. Curr. Opin. Virol. 1: 289–297.

Sabin, L.R., Hanna, S.L. & Cherry, S. 2010. Innate antiviral immunity in Drosophila. Curr. Opin. Immunol. 22: 4–9.

Scotti, P.D., Dearing, S. & Mossop, D.W. 1983. Flock House virus: a nodavirusisolated from Costelytra zealandica (White) (Coleoptera: Scarabaeidae). Arch. Virol. 75: 181–189.

Stevanovic, A. & Johnson, K.N. 2015. Infectivity of Drosophila C virus following oral delivery in Drosophila larvae. J. Gen. Virol., doi: 10.1099/vir.0.000068.

Sullivan, K., Fairn, E. & Adamo, S.A. 2016. Sickness behaviour in the cricket Gryllus texensis: Comparison with animals across phyla. Behav. Processes 128: 134–143.

Suttle, C.A. 2005. Viruses in the sea. Nature 437: 356–361.

Thomas-Orillard, M. 1984. Modifications of Mean Ovariole Number, Fresh Weight of Adult Females and Developmental Time in DROSOPHILA MELANOGASTER Induced by Drosophila C Virus. Genetics 107: 635–644.

Urquhart-Cronish, M. & Sokolowski, M.B. 2014. Gene-environment interplay in Drosophila melanogaster: Chronic nutritional deprivation in larval life affects adult fecal output. J. Insect Physiol. 69: 95–100.

Vale, P.F., Choisy, M. & Little, T.J. 2013. Host nutrition alters the variance in parasite transmission potential. Biol. Lett. 9.

Vale, P.F. & Jardine, M.D. 2015. Sex-specific behavioural symptoms of viral gut infection and Wolbachia in Drosophila melanogaster. J. Insect Physiol. 82: 28–32.

Vale, P.F. & Jardine, M.D. 2016. Viral exposure reduces the motivation to forage in female Drosophila melanogaster. Fly (Austin) 1–7.

Vale, P.F. & Little, T.J. 2012. Fecundity compensation and tolerance to a sterilizing pathogen in Daphnia. J. Evol. Biol. 25: 1888–1896.

Vézilier, J., Nicot, A., Gandon, S. & Rivero, A. 2015. Plasmodium infection brings forward mosquito oviposition. Biol. Lett. 11.

Wang, X.-H., Aliyari, R., Li, W.-X., Li, H.-W., Kim, K., Carthew, R., et al. 2006. RNA Interference Directs Innate Immunity Against Viruses in Adult Drosophila. Science 312: 452–454.

Wayne, M.L., Soundararajan, U. & Harshman, L.G. 2006. Environmental stress and reproduction in Drosophila melanogaster: starvation resistance, ovariole numbers and early age egg production. BMC Evol. Biol. 6: 57.

Webster, C.L., Waldron, F.M., Robertson, S., Crowson, D., Ferrari, G., Quintana, J.F., et al. 2015. The Discovery, Distribution, and Evolution of Viruses Associated with Drosophila melanogaster. PLOS Biol 13: e1002210.

Whitfield, A.E., Falk, B.W. & Rotenberg, D. 2015. Insect vector-mediated transmission of plant viruses. Virology 479–480: 278–289.

Wilfert, L., Long, G., Leggett, H.C., Schmid-Hempel, P., Butlin, R., Martin, S.J.M., et al. 2016. Deformed wing virus is a recent global epidemic in honeybees driven by Varroa mites. Science 351: 594–597.

Wong, R., Piper, M.D.W., Wertheim, B. & Partridge, L. 2009. Quantification of Food Intake in Drosophila. PLOS ONE 4: e6063.

Zeng, C., Du, Y., Alberico, T., Seeberger, J., Sun, X. & Zou, S. 2011. Gender-specific prandial response to dietary restriction and oxidative stress in Drosophila melanogaster. Fly (Austin) 5: 174–180.

